# Impacts of different types of florivores on flower metabolomes in the field

**DOI:** 10.64898/2026.04.30.721624

**Authors:** Selina Gaar, Caroline Müller, Thomas Dussarrat

## Abstract

Herbivory is a major biotic stress for plants, triggering the induction and modulation of diverse specialized metabolites. Such induction responses are well studied for leaves and have been shown to depend on the herbivore feeding mode. Little is known about changes in flower metabolites and chemodiversity due to florivory type. Moreover, we lack an understanding of the intraspecific variation in such responses and whether these are spatially structured.
The aromatic plant *Tanacetum vulgare,* which shows high intraspecific chemodiversity in terpene profiles, was used to examine chemotype-specific metabolic responses of flower heads to infestation by the inflorescence-infesting aphid *Macrosiphoniella tanacetaria* or the flower-feeding beetle *Olibrus* spp. under field conditions. At peak flowering, each plant received both florivory treatments on separate stems, leaving one stem herbivore-free as a control. After four days, flower heads were harvested to analyze terpenes (GC-MS) and metabolic fingerprints (LC-MS).
We found stem-specific floral metabolic responses, with florivory altering specific chemical families and their chemodiversity. Levels of a few terpenes decreased following infestation, while none increased. Untargeted analyses revealed that aphid infestation had a lower effect on flower chemistry than beetle infestation, with aphid infestation mainly causing decreases and beetle infestation predominantly leading to increases in some metabolite intensities, but little overlap across treatments and chemotypes.
Our results demonstrate that floral metabolic responses to florivory are spatially structured, florivore type-specific and shaped by plant chemotype. These findings highlight that the interplay between vascular organization, insect feeding mode, and intraspecific chemodiversity governs how flowers adjust their chemical defenses.

**One-sentence summary:** *Tanacetum vulgare* showed chemotype-specific responses to florivory by aphids (*Macrosiphoniella tanacetaria*) and beetles (*Olibrus* spp.), with aphids causing decreased and beetles increased levels of metabolic features within the same plant individuals, with little overlap in significant features across chemotypes.

## INTRODUCTION

Herbivores are one of the major drivers of chemical diversification in plants. As a result, plants have evolved diverse defense strategies, including the production of constitutive specialized metabolites that contribute to resistance against herbivores (Schoonhoven et al. 2005; Fürstenberg-Hägg et al. 2013). Beyond constitutive defenses, plants can dynamically adjust their metabolome through local or systemic induction of specialized metabolites (Schoonhoven et al. 2005; War et al. 2012). Induction responses have been mainly studied in vegetative plant organs, such as leaves and roots (Kaplan et al. 2008; van Dam 2009). In flowers, an induction of defenses against insects feeding on flowers, called florivores, may lead to trade-offs as those may also interfere with pollinator attraction (McCall and Irwin 2006; Sasidharan et al. 2023b). Despite their ecological relevance, our knowledge about florivory-induced metabolic changes in flowers remains limited (Chrétien et al. 2018; 2022).

In vegetative plant parts, herbivore-induced metabolic changes have been shown to be herbivore-specific and to depend on the insect feeding mode (Pareja et al. 2012; Rusman et al. 2019). Insects with chewing-biting mouthparts frequently induce increases in certain metabolites in leaves or roots (van Dam 2009; Mostafa et al. 2022), with responses mostly mediated via the jasmonate/ethylene pathway (Koornneef and Pieterse 2008; Erb et al. 2012). In contrast, insects with piercing-sucking mouthparts can suppress induced responses with compounds in their saliva (Walling 2008) and sometimes even lead to a reduction of specialized metabolites (Sutter and Müller 2011). Defenses against aphids are mainly mediated by the salicylic acid pathway, but some overlaps and crosstalk between signaling pathways exist (Walling 2000; Pieterse et al. 2012). In flowers, most research focused on plant responses to chewing-biting insects. For instance, larvae of *Pieris brassicae* caused rapid increases in emissions such as methanol and ethyl acetate from *Diplotaxis erucoides* flowers (Farré-Armengol et al. 2015). Likewise, flower phenolic profiles of *Brassica nigra* were distinctly altered by *P. brassicae* (Lucas-Barbosa et al. 2016). Responses in flowers to aphid infestation have been until now only rarely studied (but see Chrétien et al. 2018; 2022). For example, aphid attack of inflorescences did not cause a change in total glucosinolate concentrations in flowers of *B. nigra* (Chretien et al. 2022). Induced defense responses within a plant are often spatially structured by vascular architecture and source–sink dynamics, as the movement of signaling molecules and metabolites depends on intact transport through the vascular system (Ferrieri et al. 2015). Consequently, induction patterns can also vary among organs (Lucas-Barbosa et al. 2016) and potentially even among distinct stems within the same individual. However, beyond studies on individual target metabolites, the extent to which different florivore species attacking distinct parts of reproductive organs, i.e. flower tissues vs. phloem sap of the inflorescence stem, shape floral metabolic profiles remains unclear. Untargeted chemical analyses might provide a more holistic understanding of floral responses to various types of florivores. Besides, whether plants are capable of triggering a stem-specific response has yet received little attention.

In addition to induced responses that generate variation within a species, substantial chemical variation may already exist due to differences in constitutive chemical composition. For example, different plant species show highly different composition in certain chemical families, such as glucosinolate pattern within Brassicaceae (Fortuna et al. 2014; Kazemi-Dinan et al. 2015) or alkaloids and terpenoids in species of Asteraceae (Castells et al. 2014; Grof-Tisza et al. 2022). These so-called chemotypes (Müller et al. 2026) may show different responsiveness to environmental challenges (Grof-Tisza et al. 2022; Mamin et al. 2025). Moreover, induction responses to herbivores have often been considered by focusing on individual metabolites, whereas we know little about potential changes in the chemodiversity - the richness, relative abundance and disparity of metabolites (Hanusch et al. 2026) – of different chemical families within species. Variation in flower chemodiversity in response to biotic threats has been less explored and mostly limited to flower-pollinator interactions (Hanusch et al. 2025; Authier et al. 2026).

*Tanacetum vulgare* L. (common tansy, Asteraceae) represents an ideal system to investigate the role of plant chemodiversity in plant-insect interactions due to its high intraspecific chemodiversity in both leaf and floral terpene composition (Keskitalo et al. 2001; Sasidharan et al. 2023a). Based on their stored leaf terpene profiles, individuals can be classified into chemotypes according to their dominant terpene(s) and characteristic satellite compounds (Kleine and Müller 2011; Clancy et al. 2018). Previous studies have shown that the chemotype influenced preference behavior and the development of aphids and florivores (Jakobs and Müller 2018; Benedek et al. 2019; Sasidharan et al. 2023a). For instance, the aphid *Macrosiphoniella tanacetaria*, which is a feeding specialist on *T. vulgare*, preferred the β-thujone (B-Thu) chemotype over others, both in the laboratory and the field (Kleine and Müller 2011; Jakobs and Müller 2018). These aphids feed preferentially on the stems below inflorescences (Jakobs et al. 2019) and can therefore also be considered florivores (Rusman et al. 2019). Similarly, *Olibrus* spp. (Coleoptera: Phalacridae), which feed mainly on pollen and other tissues of Asteraceae flower heads, were influenced by chemotype differences in *T. vulgare* (Eilers et al. 2021; Sasidharan et al. 2023a). Besides, intraspecific chemodiversity not only affected insect behavior but was also associated with chemotype-specific plant responses. For instance, caterpillar feeding (*Spodoptera littoralis*) generally increased emissions of volatile organic compounds, whereas prior aphid infestation by *Metopeurum fuscoviride* modified this response in a chemotype-specific manner (Clancy et al. 2020). Wireworm herbivory (*Agriotes spp*.) induced an increase in stored root sesquiterpenoids, while aphid infestation (*M. tanacetaria*) enhanced shoot monoterpenoid emissions, with the magnitude of both responses depending on the chemotype (Newrzella et al. 2026). However, we still lack an understanding of whether flower head chemodiversity is influenced by distinct florivore types and whether these responses are chemotype-specific.

Here, we explored the consequences of florivory by insects with different feeding modes on the chemical response of *T. vulgare* flower heads in a field setting, focusing on impacts on chemodiversity of chemical families as well as changes in individual metabolites. More specifically, we asked (i) whether a plant can orchestrate a stem-specific response to two distinct types of florivory (*M. tanacetaria* and *Olibrus* spp.) and (ii) whether this chemical response is insect species- and plant chemotype-dependent. Stems of flowering plants were either kept insect-free as a control or infested with aphids or beetles. Flower heads were collected after four days and subject to targeted (terpene profile analysis) and untargeted metabolomics. We expected insect infestation to cause only minor changes in stored terpene profiles, with overall variation being dominated by differences specific to the plant’s chemotype. Furthermore, depending on insect species and plant chemotype, we predicted plants to increase the production of several floral metabolites, with a lower responsiveness to aphid infestation.

## MATERIALS AND METHODS

### Plant material

Plants of *Tanacetum vulgare* were grown in an outdoor common garden near Bielefeld University, Germany (52°03′39.43″N, 8°49′46.66″E; elevation 142 m; for details see Ziaja and Müller (2023)). Briefly, plants originated from seeds collected in 2019 at four sites in Bielefeld. Based on leaf monoterpene profiles, plants of five distinct chemotypes were selected and planted individually in polyvinyl chloride tubes (diameter 16 cm, height 30 cm, inserted 25 cm into the soil) in a common garden in May 2020 in several blocks (300 plants in total, set-up in Fig. S1). The garden was managed uniformly across years, including regular vegetation removal and annual winter cutting. Due to climatic events, only thirty plants flowered four years later in 2024, which were used for the present experiment and belonged to three chemotypes, being either dominated (> 50% of total terpene concentration) by α- and β-thujone (called “AB-Thu” in the following, *n* = 9), β-thujone (“B-Thu”; *n* = 11) or artemisia ketone (“Keto”, *n* = 10). In each tube, several separate stems had grown from the rhizome.

### Insect collection and maintenance

*Macrosiphoniella tanacetaria* aphids were collected in May 2024 from *T. vulgare* plants in Bielefeld (52°02′14.3″ N, 8°29′23.6″ E) and reared subsequently on *T. vulgare* plants grown from seeds (Blauetikett Bornträger GmbH, Germany) to obtain a random mixture of different chemotypes. Aphids were kept on plants in a tent (60 × 60 × 60 cm, Bugdorm BD2F120; MegaView Science, Taiwan) in a climate chamber under a 16-hour light/8-hour dark photoperiod at 22 °C/18 °C with 60 –70 % relative humidity. During this period, the aphids reproduced and were kept in the laboratory for around two months before use in the experiment.

Around 200 adult *Olibrus* spp. beetles (probably *Olibrus aeneus*) were collected from June to July 2024 from flower heads of wild chamomile (*Matricaria* spp., *Anthemis* spp. or *Tripleurospermum* spp., Asteraceae) in Bielefeld (52°02’51.4”N, 8°29’51.5”E) and Salzkotten, Germany (51°41’34.3”N, 8°33’26.0”E). Around 50 beetles were kept in plastic boxes (22 x 13 x 12 cm) lined with paper tissue in the same climate chamber as the aphids. Beetles were fed with fresh flower heads of *T. vulgare* from a mixture of five different chemotypes kept in a greenhouse, as well as flower heads collected from wild *T. vulgare*. Although mating behavior was observed, no oviposition or larval development occurred under laboratory conditions. Therefore, only field-collected adult beetles that had been in the laboratory for at least two weeks were used in the experiment.

### Field experiment and sampling

At peak flowering, each *T. vulgare* plant received three treatments: infestation with *M. tanacetaria*, infestation with *Olibrus* spp. and an uninfested control (Fig. 1; Table S1). For each treatment, flower heads on separate stems of the same plant, emerging from the ground and being connected by the rhizome, were enclosed in mesh bags (11 × 11 cm), resulting in three bags per plant. Depending on plant architecture and size, each bag contained 5–16 flower heads. In four plants (one AB-Thu, one B-Thu, two Keto) having less than three distinct stems, flower heads on different branches expanding from the same rosette were used instead. Insects were introduced by cutting a small hole in one corner of the mesh bag, inserting the insects and sealing the opening with a paper clip. Control bags were handled identically but received no insects. Herbivore densities were standardized across plants, with two apterous adult aphids per flower head and one adult beetle per three flower heads, respectively. These infestation levels were based on observations of natural herbivore densities on wild *T. vulgare* plants. The infestation period lasted four days, after which the insects were removed. Flower heads were then harvested, immediately flash-frozen in liquid nitrogen and stored at −70 °C. Infestation and harvest were conducted once per plant at peak flowering between 10 am and 1.30 pm from late July to late August. Although climatic conditions varied among sampling dates, we prioritized maintaining a consistent plant developmental stage (peak flowering) across all experimental runs. Therefore, significant treatment effects reflect robust induction responses that are reproducible across varying climatic conditions. Prior to extraction, frozen flower heads were freeze-dried for 48 h and ground to a fine powder. The experimental design and workflow are summarized schematically in Fig. 1.

**Fig. 1.**
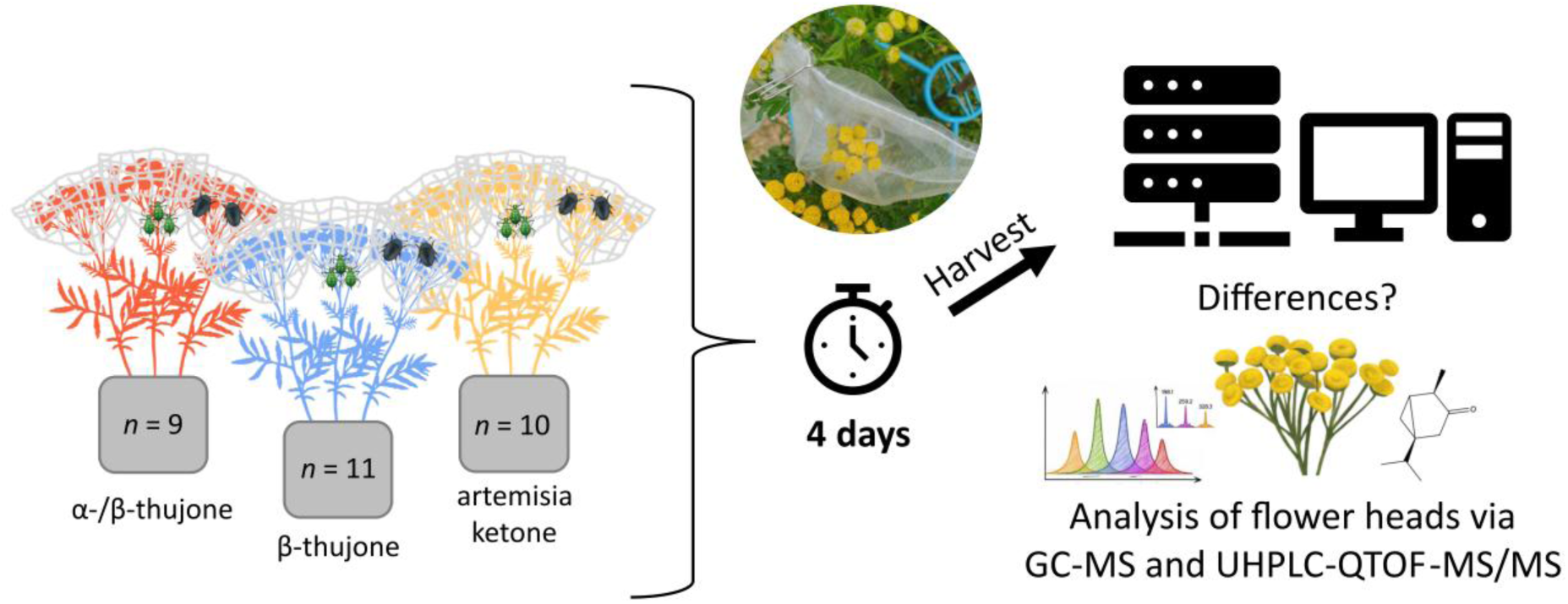
Experimental setup for analyzing differences in floral responses of *Tanacetum vulgare* to infestation by herbivorous insects. Plants of three chemotypes (α-/β-thujone: *n* = 9, β-thujone: *n* = 11, artemisia ketone: *n* = 10) were used to assess chemotype-specific responses. Each plant received three treatments on different stems. Depending on the number of available flower heads per bouquet, between five and 16 flower heads were enclosed in mesh bags. The number of insects per bag was standardized to the number of flower heads: two aphids (*Macrosiphoniella tanacetaria*) were added per flower head and one beetle (*Olibrus* spp.) per three flower heads. Control bouquets received empty mesh bags. Insects were allowed to feed for four days, after which flower heads were harvested and analyzed for terpenoids (GC–MS) and metabolic fingerprints (UHPLC–QTOF–MS/MS).

### GC-MS analysis of stored terpenes

Stored terpenes were analyzed following established protocols with minor modifications (Eilers et al. 2021; Ziaja and Müller 2025). For each sample, 10 ± 2 mg of dried, ground flower head material was weighed into FastPrep tubes and extracted with 1 mL *n*-heptane (HPLC grade, Roth, Germany) containing 10 ng µL⁻¹ 1-bromodecane (GC-MS grade, Sigma Aldrich, Steinheim, Germany) as an internal standard. Approximately 150 µL of ceramic beads were added for homogenization. Samples were processed in a FastPrep-24^TM^ 5G homogenizer (MP Biomedicals, Eschwege, Germany) for 30 s at 9 m s⁻¹, followed by sonication in an ultrasonic bath for 5 min at room temperature. Samples were then centrifuged at 13,200 rpm for 10 min at 20 °C and the supernatants were used for GC–MS analysis. Extraction blanks containing only the internal standard were measured every 20 samples to detect potential contamination.

Samples were analyzed by GC–MS (GC 2010plus – MS QP2020, Shimadzu, Kyoto, Japan) operated in electron impact ionization mode at 70 eV. Compounds were separated on a semi-polar VF-5 MS capillary column (30 m length, 0.25 mm ID, 10 m guard column; Agilent Technologies, Santa Clara, CA, USA) using helium as carrier gas at a constant flow of 1.5 mL min⁻¹. Samples were injected at 230 °C in split mode (1:10). The oven temperature program started at 50 °C, held for 5 min, increased to 250 °C at 10 °C min⁻¹, followed by a ramp to 280 °C at 30 °C min⁻¹ and a final hold of 3 min. The MS interface was set to 250 °C and the ion source to 230 °C. Mass spectra were acquired in quadrupole MS mode with the following scan settings: from 8.2 to 19 min, scan speed 2000 and *m*/*z* range 30–400; from 19 to 29 min, scan speed 625 and *m*/*z* range 30–500. An alkane standard mixture (C7–C40; Sigma-Aldrich) was analyzed with the same GC–MS method to calculate retention indices (RI) according to van den Dool and Kratz (1963). Chromatograms were analyzed using LabSolutions GC-MS Postrun Analysis v4.45 (Shimadzu).

Terpenes were identified by comparison of their RI and mass spectra reported in Adams (2007) and entries from the mass spectral library of the NIST 2014 and the FFNSC 3 (Mondello 2015), as previously described in Ziaja and Müller (2025). Compounds were considered identified if MS spectra matched reference spectra and if RI deviations did not exceed 1.5% of reported values (Adams 2007). Other features were annotated as “unknown monoterpene or sesquiterpene”. Compounds were quantified based on peak areas from the total ion chromatogram, normalized to the internal standard. The mean peak areas of the blanks were subtracted from each sample and values were subsequently normalized to the sample dry mass. Compounds detected in fewer than two samples were removed from the table, yielding a total of 102 putatively annotated terpenes (Tables S2, S3).

### LC-MS analysis of metabolic fingerprints

Metabolic fingerprints were analyzed following established protocols with minor modifications (Schweiger et al. 2021; Dussarrat et al. 2023). For each sample, 10 ± 2 mg of dried, ground flower material was extracted with 500 µL of 90 % methanol (v/v) containing hydrocortisone (10 mg L⁻¹) as an internal standard (Sigma-Aldrich). Samples were vortexed and subsequently sonicated for 20 min in an ice-cooled ultrasonic bath. Extracts were then centrifuged for 10 min at 6 °C and 13,200 rpm. The supernatant was filtered through a 0.2 µm syringe filter (Phenomenex, Torrance, CA, USA) and used for LC–MS analysis. Extraction blanks containing only the internal standard were prepared in parallel to monitor potential contamination. In addition, quality control samples were prepared by pooling aliquots of all extracts.

Samples were analyzed by LC-MS (UHPLC-QTOF-MS/MS; UHPLC: Dionex UltiMate 3000, Thermo Fisher Scientific, San José, CA, USA; QTOF: compact, Bruker Daltonics, Bremen), equipped with a Kinetex XB-C18 column (150×2.1 mm, 1.7 µm, with guard column; Phenomenex), following Schweiger et al. (2021) with minor modifications. Compounds were separated at a flow rate of 0.5 mL min⁻¹ at 45 °C using Millipore water containing 0.1 % formic acid as eluent A and acetonitrile containing 0.1 % formic acid as eluent B. The gradient started at 2 % B, increased to 30 % B over 20 min, followed by an increase to 75 % B within 9 min and ended with column cleaning and re-equilibration. Blanks were injected every 20 samples and quality controls every 15 samples to ensure measurement quality. Measurements were performed in negative electrospray ionization mode with an *m*/*z* range of 50–1300 and a spectra acquisition rate of 6 Hz. Further instrument settings were applied as described in Schweiger et al. (2021). Signals acquired in MS mode were used for quantitative analyses. Additional fragmentation spectra were acquired in AutoMS/MS mode, using a separate chromatographic trace for features that exceeded a predefined intensity threshold. Nitrogen was used as the collision gas. Prior to each sample run, sodium formate calibration solution was introduced into the system for recalibration of the *m*/*z* axis. For selected samples, MS/MS analyses were performed with MRM to fragment selected ions at 8 Hz. The isolation widths and collision energies were defined according to the *m*/*z* of the precursor ions.

Feature detection, alignment and grouping were performed using the T-ReX 3D algorithm implemented in MetaboScape (v. 2021b, Bruker Daltonics). Raw LC–MS data were processed to detect features defined by a specific *m*/*z* at a given retention time (RT). Features within an RT window of 0.3–29 min were considered for further analysis. Feature detection was performed using an intensity threshold of 1000 counts, a minimum peak length of 15 spectra (for MRM measurements 7 spectra) and an EIC correlation threshold of 0.8. Only the features that were present in at least two samples were retained. Feature intensities were normalized by the internal standard (hydrocortisone [M+HCOOH-H]⁻) and sample dry weight. Mean normalized blank intensities were then subtracted from each corresponding feature and negative values were set to zero. During LC-MS preprocessing, six samples were excluded due to large retention time shifts. The datasets are available in Table S4 and S5.

Chemical class annotation was performed using SIRIUS v6.1.0 (Dührkop et al. 2019) as previously described in Authier et al. (2026). While all parameters are available in Table S6, the metabolite structure and chemical class predictions were performed using CSI:FingerID (Hoffmann et al. 2022) and CANOPUS (Dührkop et al. 2021). For each chemical feature, the pathway, superclass and class (Natural Products Classifier ontology) were assigned if the confidence score was ≥ 0.8 (Table S7). For each chemical family containing at least ten features, chemodiversity indices (richness, Shannon index and inverse Simpson index) were calculated. To consider disparity, the cosine similarity score between MS/MS spectra was assessed via the Global Natural Product Social Molecular Networking web platform (GNPS) (Wang et al. 2016) using feature-based molecular networking (Nothias et al. 2020). Cosine similarity scores were converted to dissimilarity values (1 – cosine score) to generate dissimilarity matrices for all chemical families. Functional Hill diversity (FHD) indices were then calculated using the chemodiv package (Petrén et al. 2023). In total, 76 chemodiversity indices were obtained (Table S8). All chemical datasets are available in the supplemental material and were deposited online (see Data Availability section).

### Data analysis

All data analyses was performed using the tidyverse collection of packages, including dplyr, tidyr, stringr, tibble, readr, purrr and ggplot2 (Wickham et al. 2019) on R v4.5.2 (R Core Team 2025). Multivariate analyses were conducted and additional figures were developed using the packages car (Fox and Weisberg 2019), ARTool (Kay et al. 2025), mixOmics (Rohart et al. 2017), ComplexUpset (Krassowski et al. 2022), vegan (Oksanen et al. 2025) and pheatmap (Kolde 2019).

All analyses accounted for the paired experimental design in which each plant received all three treatments. The effects of florivory treatment on terpene chemodiversity (richness, Shannon index and inverse Simpson index) and how these effects depend on plant chemotype were tested. To test the influence of treatment, chemotype and their interaction on the indices, non-parametric factorial analyses based on the Aligned Rank Transform (ART) approach were performed. *P* < 0.05 was considered as significant. To explore the variance in the chemical dataset, unsupervised analyses (standard and multilevel principal component analysis, PCA) and multilevel partial least squares discriminant analysis (PLS-DA) were performed on normalized GC-MS data [relative values, cube root transformation and Pareto scaling performed in MetaboAnalyst v6.0 (Pang et al. 2024)]. To identify compounds responding significantly to florivory and chemotype, volcano plots based on absolute data were generated, calculating Wilcoxon tests and fold changes (FC) for treatments *versus* controls, both across all samples and within each chemotype. *P*-values from Wilcoxon tests were adjusted for multiple testing using the false discovery rate (FDR) method according to Benjamini and Hochberg (1995). As no terpenes remained significant after FDR correction, those with *P* < 0.05 in the uncorrected Wilcoxon tests were visualized in a heatmap based on normalized data.

Similarly, the effects of treatment, chemotype and their interaction on calculated LC-MS chemodiversity indices across chemical families were analyzed using the ART approach. To account for multiple testing, *p*-values from the ART models were adjusted using FDR correction according to Benjamini and Hochberg (1995). Variation in metabolic fingerprints was assessed using unsupervised PCA and multilevel PLS-DA on normalized LC-MS data (median normalization, cube root transformation and Pareto scaling performed in MetaboAnalyst). As the multilevel approach requires paired data, one sample without a corresponding pair (due to prior exclusions caused by retention time shifts) was removed. Differences in the variability of metabolic profiles among treatments were assessed using the *vegan* package via a permutation-based test for homogeneity of multivariate dispersions (PERMDISP) based on Bray–Curtis dissimilarities with 999 permutations. To identify insect species- and chemotype-specific modulations (*P* < 0.05), volcano plots were generated using the same approach as for the GC-MS data. As paired comparisons were required, five samples without a corresponding pair (due to prior exclusions caused by retention time shifts) were removed. Significant features identified by Wilcoxon tests in each group were visualized in an UpSet plot. To further characterize the putatively biologically relevant changes, features that met both the Wilcoxon significance criterion (before FDR correction) and the FC threshold were annotated and classified according to the Metabolomics Standards Initiative (MSI) confidence levels (Sumner et al. 2007; Schymanski et al. 2014).

## RESULTS

### Consequences of florivory and chemotype on the terpene profile

A total of 102 terpenes were detected in flower heads by GC-MS. Chemotypes differed in richness, Shannon and inverse Simpson diversity (Tables S9, S10). Florivory treatment had no significant effect on any of the terpene chemodiversity indices (richness, Shannon and inverse Simpson diversity) and no significant interaction between florivory treatment and chemotype was detected (Tables S9, S10).

Standard PCA revealed chemotype-related structuring of terpene profiles (Fig. S2A). Samples of the Keto chemotype formed a distinct cluster, whereas samples of the AB-Thu and the B-Thu chemotype largely overlapped in the multivariate space, with no separation among florivory treatments (Fig. S2A). Multilevel PCA, focusing on within-plant variation, showed no distinct clustering of samples according to insect treatment, indicating only minor florivory-induced changes in overall terpene composition (Fig. S2B). As the experiment was conducted on different stems of the same plant individual, observed variation among treatments reflects within-plant responses rather than differences among independent individuals. Similarly, multilevel PLS-DA revealed no clear separation of terpene profiles among florivory treatments. However, control samples tended to cluster separately from insect-treated samples, whereas beetle- and aphid-treated samples largely overlapped (Fig. 2A). The multilevel PLS-DA revealed substantial differences in classification performance among treatments. Samples of the beetle treatment showed the highest error rates across both components, whereas control and aphid samples were classified with comparatively lower error rates, particularly on the first component.

**Fig. 2.**
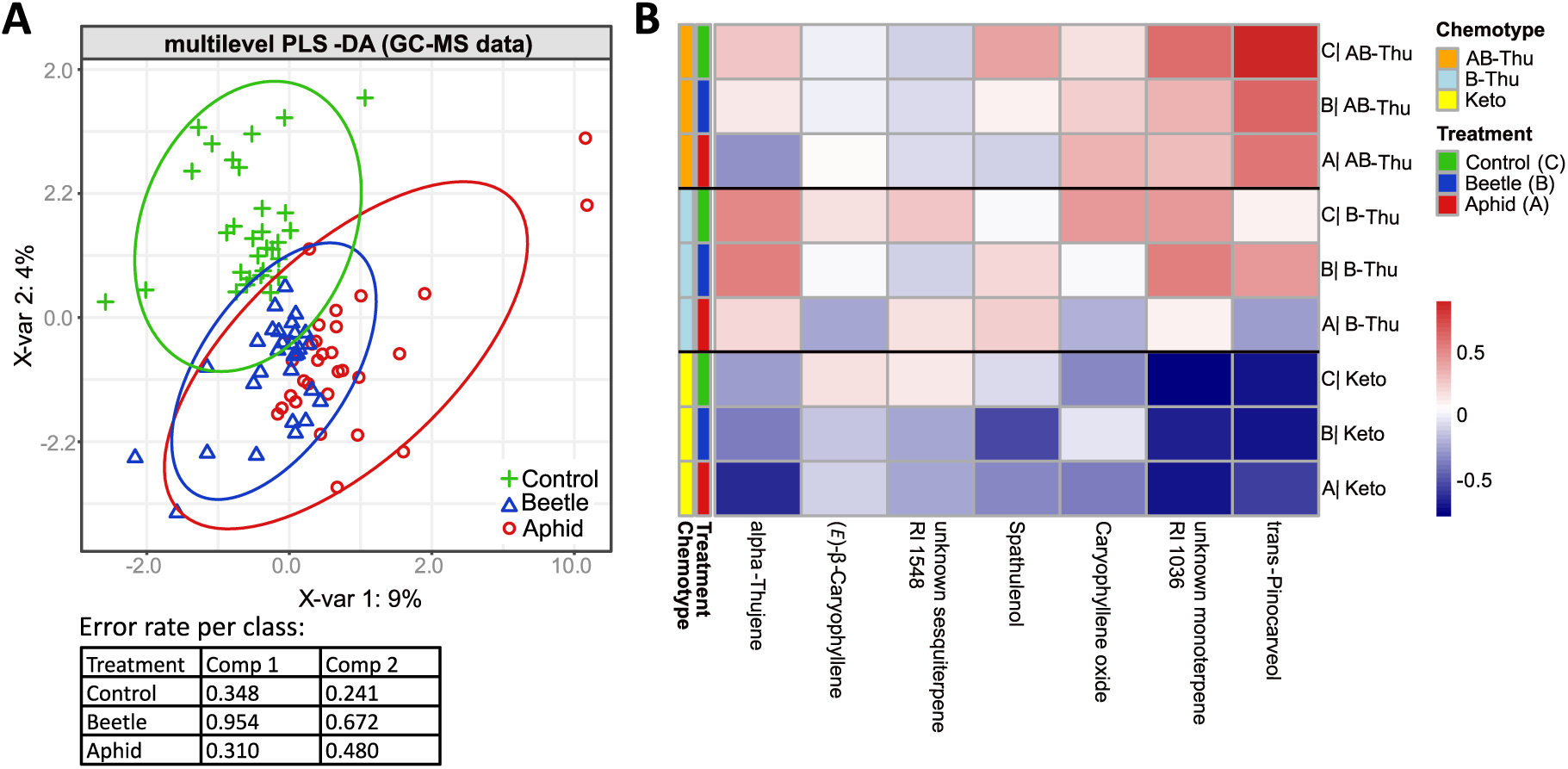
Stored terpene profiles of *Tanacetum vulgare* flower heads in response to insect infestation. A. Multilevel PLS-DA score plot showing the separation of samples between the three treatment groups (Control, Beetle, Aphid) in a paired dataset, where each plant received all three treatments on different stems. Analysis was performed on normalized GC-MS data (102 terpenes). Symbols and colors indicate the treatments: green crosses represent uninfested flower head control samples, red circles represent samples infested with aphids (*Macrosiphoniella tanacetaria*) and blue triangles represent samples infested with beetles (*Olibrus* spp.). Ellipses indicate the 95 % confidence intervals for each treatment group. B. Heatmap of the group mean values of normalized terpene abundances of seven terpenes that differed significantly between treatments in Wilcoxon tests (insect treatment [Aphid or Beetle] *versus* Control, *P* < 0.05), analyzed across all samples and chemotype-specifically (Keto: artemisia ketone chemotype; B-Thu: β-thujone chemotype; AB-Thu: α-/β-thujone chemotype). Rows correspond to treatment groups within each chemotype, columns correspond to the seven terpenes and colors represent scaled abundances (red = high, blue = low).

### Florivore- and chemotype-specific responses of individual terpenes

Although florivory did not reshape the overall terpene profile, it influenced the abundance of particular terpenes (Fig. S3). We first explored the terpene response to aphid and beetle florivory across all plants, without considering chemotypes (Fig. S3A and S3B). Aphid treatment had no significant effect on any terpene (Fig. S3A), whereas beetle florivory significantly reduced the levels of two sesquiterpenes, (*E*)-β-caryophyllene and an unknown sesquiterpene (RI 1548) (Fig. S3B). Next, we tested the influence separately per terpene chemotype (Fig. S3C to S3H). In the B-Thu chemotype, one terpene (caryophyllene oxide) showed a significant decrease under aphid treatment (Fig. S3E). In addition, three terpenes responded significantly to beetle infestation and one terpene to aphid infestation (*P* < 0.05, Wilcoxon test), but did not meet the fold change threshold (FC < 0.5 or FC > 2) (Fig. S3C-D, F-H, Table S11). Overall, these seven significant terpenes also showed pronounced variation in abundance and responsiveness to herbivores between chemotypes (Fig. 2B). Importantly, no significant increases in terpene levels were observed in any comparison. No compound responded significantly after the FDR correction.

### Responses in chemodiversity indices from multiple chemical families and individual metabolites to insect infestation across chemotypes

A total of 3742 features were detected in flower heads by LC-MS. Nineteen chemical families met the criterion of containing at least ten features and were included in the chemodiversity analyses (richness, Shannon index, inverse Simpson index and FHD; Table S8). Terpene chemotype influenced five chemical indices, primarily affecting monoterpenoids, whereas treatment mainly impacted phenolic acids across five indices. The chemotype × treatment interaction affected three indices, all associated with fatty acyl glycosides. Significant effects were most frequently observed for the inverse Simpson index (five cases), followed by richness and the Shannon index (three cases each) and the FHD (two cases) of specific chemical families. Following FDR correction, only the effect of chemotype on monoterpene richness remained significant (Table S12).

Multilevel PCA, focusing on within-plant variation, showed no distinct clustering of samples by treatment, consistent with only minor changes in overall metabolic profiles (Fig. S4). The main variation captured by PC1 likely reflects residual within-plant variation among different stems of a plant individual rather than treatment-specific effects. Similarly, multilevel PLS-DA analysis revealed no clear separation of metabolic fingerprints among florivory treatments. However, control samples tended to cluster separately from beetle-treated samples, whereas aphid-treated samples largely overlapped with the other two groups (Fig. 3). The multilevel PLS-DA revealed pronounced differences in classification performance among treatments. Aphid samples showed very high error rates on both components, indicating poor discrimination from the other groups. In contrast, control and beetle samples exhibited substantially lower error rates, with beetle samples being particularly well classified on the second component. While the first component moderately separated control and beetle samples, aphid-induced metabolic fingerprints were not distinctly captured by the model. In line with the similar dispersion of the treatment groups observed in the multilevel PLS-DA ordination space, the multivariate dispersion of the metabolic fingerprints did not differ between treatments (PERMDISP, *p* > 0.05, Table S13), indicating a limited influence of florivory treatment on the overall flower metabolome.

**Fig. 3.**
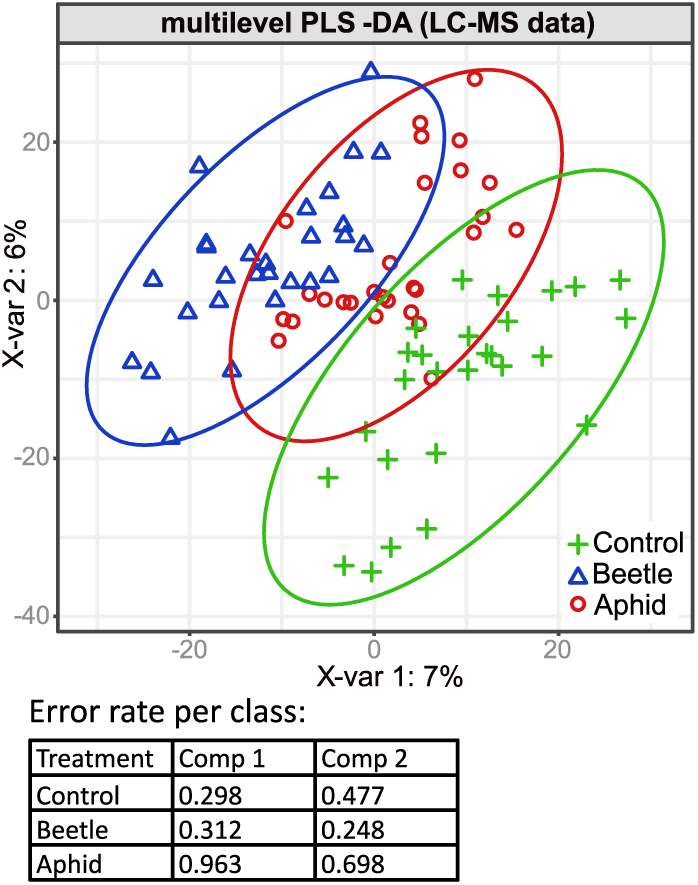
Metabolic fingerprints of *Tanacetum vulgare* flower heads in response to insect infestation. Multilevel PLS-DA score plot showing the separation of samples between three treatment groups (Control, Beetle, Aphid) in a paired dataset, where each plant received all three treatments on different stems. Analysis was performed on normalized data of LC-MS metabolic fingerprints (3731 features). Symbols and colors indicate the treatments: green crosses represent uninfested flower head control samples, red circles represent samples infested with aphids (*Macrosiphoniella tanacetaria*) and blue triangles represent samples infested with beetles (*Olibrus* spp.). Ellipses indicate the 95 % confidence intervals for each treatment group.

### Aphid and beetle florivory trigger distinct floral chemical traits, with a chemotype-specific response

At the chemical feature level, statistical analyses revealed distinct metabolic responses to aphid and beetle infestation, as well as differences among chemotypes (Figs. 4 and 5). Without considering chemotype differences, beetle infestation resulted in a higher number of responding features than aphid infestation (Fig. 4). Moreover, aphid-infested flower heads showed a predominance of features with decreased levels relative to controls (Fig. 4A), whereas beetle infestation was characterized by a predominance of features with increased levels (Fig. 4B). This pattern remained consistent when chemotypes were analyzed separately, with beetle infestation generally associated with more features increased and aphid infestation with more features decreased in level, despite a reduced number of significant features (Fig. S5). However, none of the features remained significant after applying the Benjamini–Hochberg FDR correction. A detailed summary of the results for each individual feature is given in Table S14.

**Fig. 4.**
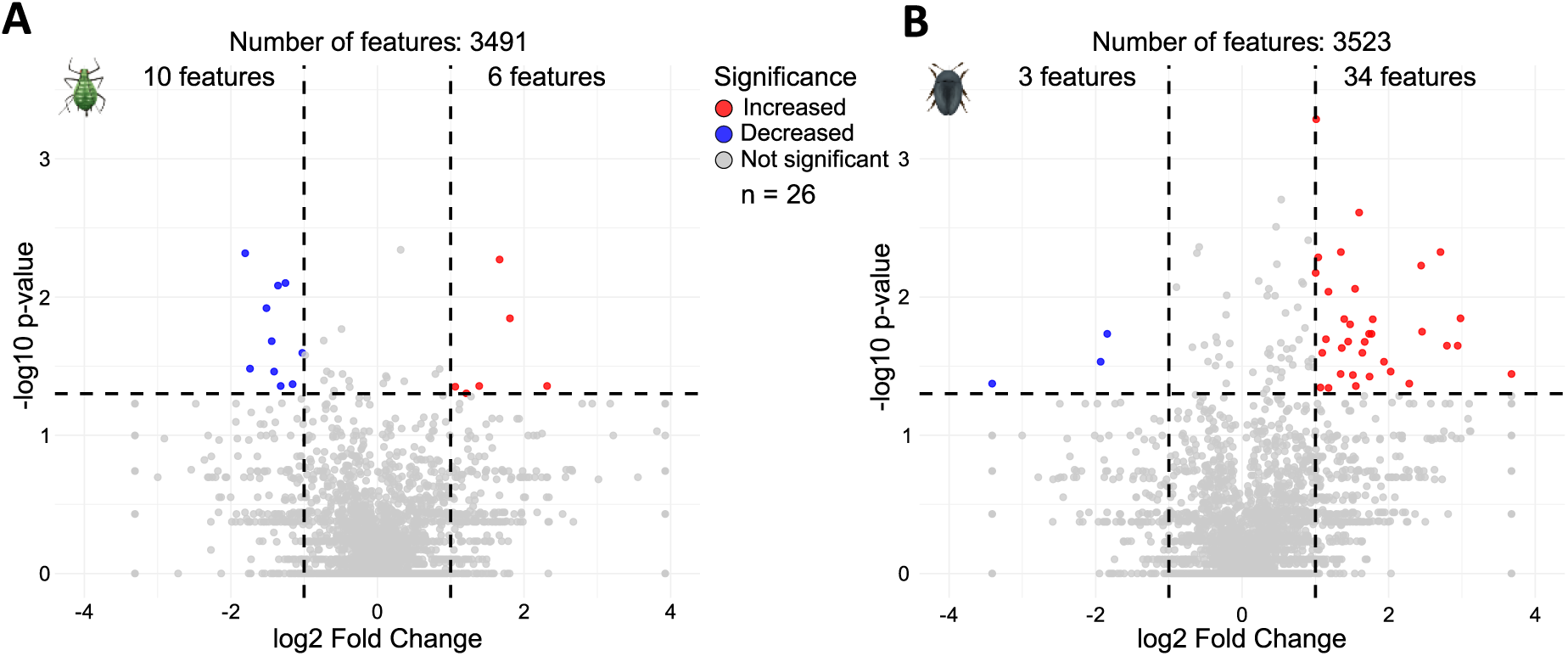
Species-specific metabolic responses of *Tanacetum vulgare* flower heads to aphid and beetle infestation. Volcano plots showing metabolic features significantly modulated by insect infestation, displayed as circles outside the cutoff lines (Wilcoxon test, *P* < 0.05; fold change < 0.5 for decreases and > 2 for increases) based on absolute LC-MS data. Blue circles indicate features with reduced levels in infested flower heads compared to controls, while red circles indicate increased levels. For features detected exclusively in one treatment group, fold changes were set to the maximum observed value for decreases or increases within the respective comparison. Sample sizes were *n* = 26 plant pairs for aphids-control as well as for beetles-control. The number of metabolites shown at the top center represents the total number of features detected for each treatment comparison, while the numbers at the top left and right indicate the counts of significantly decreased and increased features, respectively. A. Aphid *versus* control B. Beetle *versus* control.

**Fig. 5.**
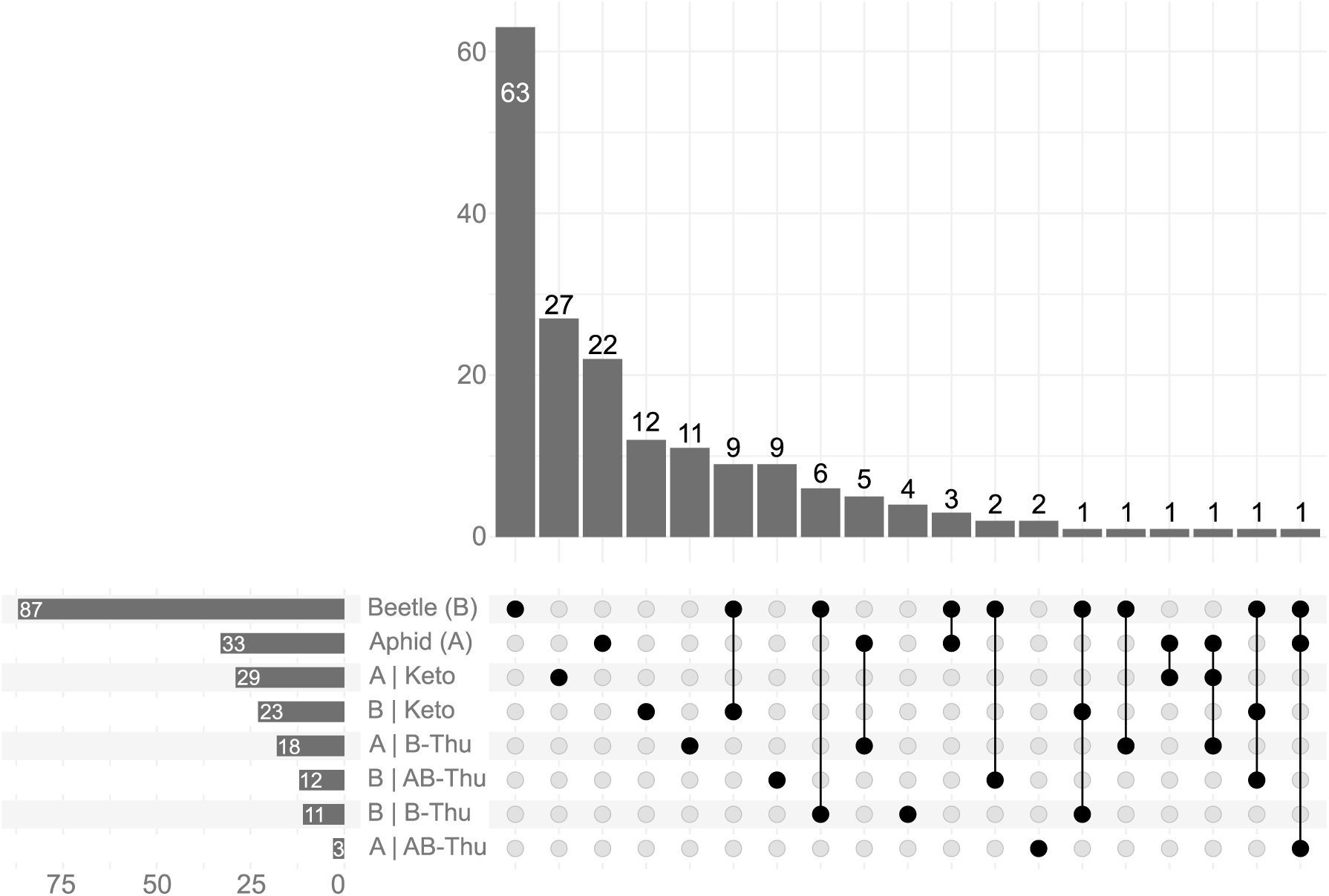
Chemotype- and herbivory-dependent variation in metabolic responses of *Tanacetum vulgare* flower heads. Upset plot showing the number of significant features identified by Wilcoxon tests (insect treatment [A = Aphid; B = Beetle] *versus* Control, *P* < 0.05), conducted for all chemotypes combined as well as for each chemotype individually (Keto: artemisia ketone chemotype; B-Thu: β-thujone chemotype; AB-Thu: α-/β-thujone chemotype). In total, 3731 features were detected across all samples. Bars on the left indicate the total number of significant features within each group, while bars on the top represent intersections between groups, illustrating shared and unique metabolic responses to insect infestation.

Next, we explored the overlap and treatment specificity of significant features between aphid- and beetle-infested samples and assessed how these responses varied across chemotypes (Fig. 5). The overlap of significant chemical features between insect treatments and among chemotypes was minimal, indicating largely distinct floral chemical responses. Of the 181 significant features identified in aphid- and beetle-infested flower heads and chemotypes, only five were shared between the two insect treatments. Similarly, flower head responses to aphid and beetle infestation involved 74 and 112 chemical features, of which 65% and 38% were chemotype-specific, respectively (Fig. 5).

Subsequently, we putatively annotated the 79 features (i.e. 53 from chemotype-independent analyses and 26 additional features from chemotype-specific analyses) that met both the Wilcoxon significance criterion and the FC threshold (Table S15, Fig. 6). Pathway classification indicated that most responsive features, irrespective of insect species, were associated with the shikimate and phenylpropanoid pathways and generally showed increased levels in both treatments (Fig. 6). Despite this convergence at the pathway level, only two individual features were significantly affected by both insect treatments and both increased. One was assigned to the terpenoid pathway and the other was an unknown feature. In aphid-infested samples, additional affected features represented fatty acid, terpenoid and alkaloid pathways. Besides, aphid infestation was more frequently associated with decreased levels of features across pathways, with the shikimate and phenylpropanoid pathways representing a notable exception, as certain compounds such as eugenol showed increased levels irrespective of chemotype. By contrast, beetle infestation was primarily characterized by increases in feature levels, with exceptions such as the decrease of the phenolic compound piceol in the B-Thu chemotype. Additional features were assigned to fatty acid, terpenoid, carbohydrate, amino acid and peptide pathways. Phenylpropanoids (C6–C3) were particularly prominent. Within this group, cinnamic acids and their derivatives represented the dominant class (Table S15). Furthermore, one feature was exclusively detected in beetle-infested samples and was absent in controls. However, this feature does not provide reliable structural identification. Its absence in the control group may also indicate concentrations below the limit of detection rather than a true absence.

**Fig. 6.**
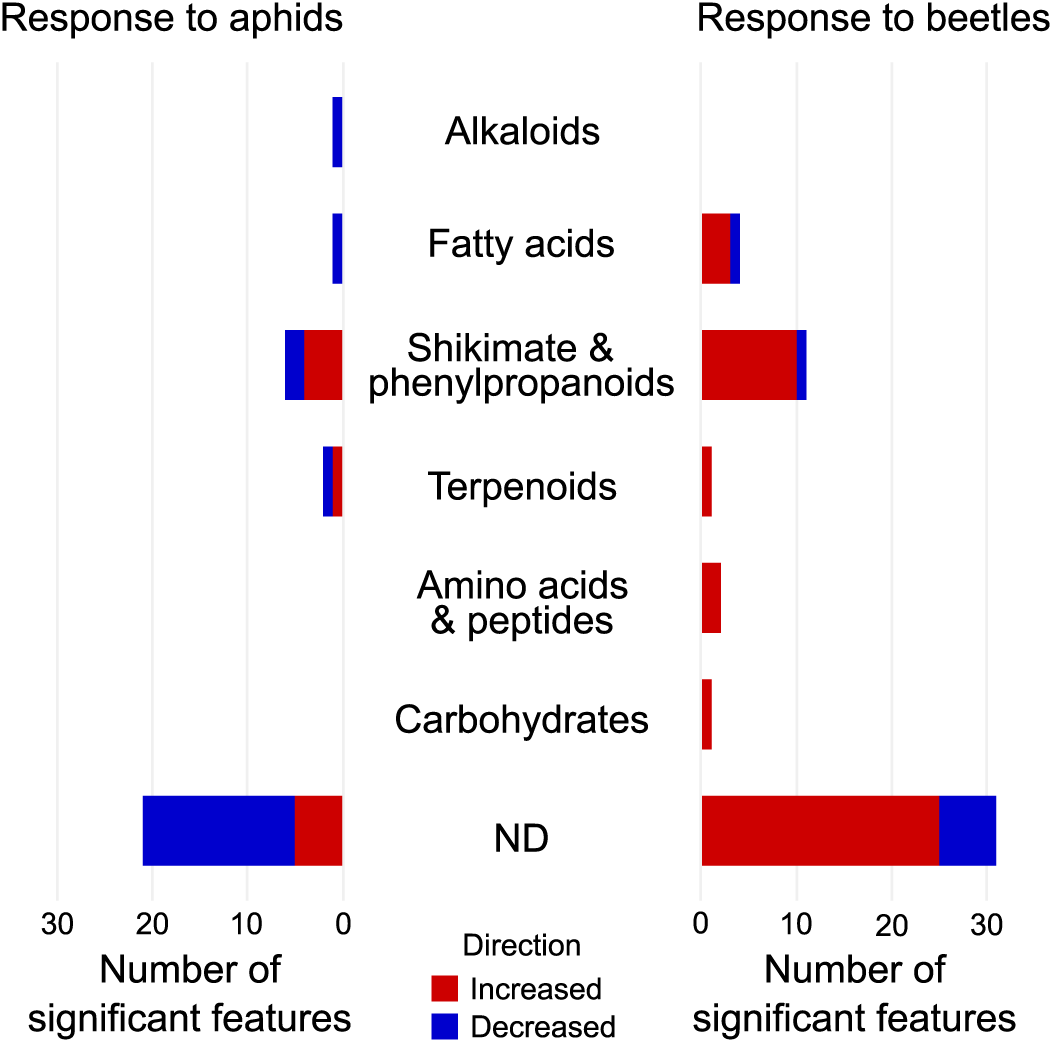
Pathway classification of significant features induced in *Tanacetum vulgare* flower heads by aphid and beetle infestation. Bar plots showing the number of significant features based on volcano plot analyses conducted for all chemotypes combined as well as for each chemotype individually. In total, 79 features responded significantly (Wilcoxon test, *P* < 0.05; fold change < 0.5 for decreases and > 2 for increases). Bars on the left indicate the number of features responding to aphid infestation, whereas bars on the right indicate the number responding to beetle infestation across the different chemical pathways. Red indicates the proportion of features increased in level and blue the proportion decreased. ND = not determined. Two features were shared between both insect treatments, with one assigned to the terpenoid pathway and the other classified as ND; both increased in response to aphid and beetle infestation.

## DISCUSSION

Plant defense mechanisms and chemical diversity have fascinated chemical ecologists for decades. However, florivory-induced metabolic changes in flowers received only limited attention compared to vegetative organs, especially at the chemodiversity level. Besides, the extent to which plants are capable of orchestrating stem-specific chemical responses to different types of florivores and how these responses are modulated by intraspecific chemodiversity, such as distinct terpene chemotypes, remains poorly explored. Here, we investigated the impact of florivore type and chemotype on stem-specific chemical responses of *Tanacetum vulgare* flower heads in a field setting, at the level of chemodiversity indices of different chemical families and at the individual metabolite level. Our findings support stem-specific floral metabolic responses in *T. vulgare*. Both the floral terpene profile and the untargeted metabolome were influenced by insect species and terpene chemotype. However, these effects were comparatively weak at the level of stored terpenes and more pronounced in the untargeted metabolomic analyses.

### Flower heads exhibit stem-specific chemical responses to florivory

Our results indicate that floral metabolic responses varied between differently treated stems of the same plant, suggesting a substantial degree of stem-specific regulation, with flowers responding locally to florivory on the affected stem rather than exhibiting uniform systemic induction across the plant. Previous findings in *Arabidopsis thaliana* revealed that systemic responses depend on vascular connectivity, with the strongest induction occurring in leaves within the same orthostichy as the leaves damaged by leaf chewers (Ferrieri et al. 2015). Similarly, herbivory by *Spodoptera littoralis* increased resistance in developing leaves throughout cotton plants, although local leaves often responded more strongly than distal ones, generating substantial intra-plant variation (Anderson and Agrell 2005). Such findings emphasize that plant defenses are organized along vascular connections and architecture rather than being uniformly distributed. However, vascular connectivity does not necessarily imply full physiological integration. Although individual stems of *T. vulgare* were connected via a shared rhizome at the vascular level, our results indicate that floral metabolic responses remained largely stem-specific. This suggests that, despite rhizome-mediated integration, systemic defense signaling between stems is spatially constrained, pointing to a high degree of modular autonomy. Such stem-specific induction may represent an adaptive balance between protection and maintenance of other essential functions, particularly in reproductive tissues. Restricting chemical responses to attacked stems may minimize costs and reduce potential trade-offs with pollinator attraction (Strauss et al. 2002; Sasidharan et al. 2023b). Stem-specific regulation may therefore enable plants to fine-tune floral defense while preserving overall reproductive performance.

### Florivory induces selective rather than global metabolic responses in flower heads

Florivory did not alter the overall terpene profile of *T. vulgare* flower heads but affected the abundance of specific terpenes, with only a few significant decreases in sesquiterpenes and no increases observed. This supports our expectation that stored terpenes exhibit limited inducibility. Previous studies have shown that stored terpenes tend to exhibit weaker or less dynamic responses than volatile emissions in some systems. A comparable pattern was reported by Newrzella et al. (2026), who found that infestation of *T. vulgare* leaves by *Macrosiphoniella tanacetaria* aphids reduced the concentration of stored shoot sesquiterpenes while leaving stored monoterpene pools unaffected. In their study, changes in the emission of volatile terpenes were observed in both terpene classes. In wild cotton, where both stored and emitted volatiles increased after herbivory by *Spodoptera exigua*, chemotype-defining terpenes showed weak inducibility and the constitutive profile largely persisted (Mamin et al. 2025), indicating that chemotype-specific differences often dominate overall terpene variation.

Beyond terpenes, our untargeted analysis highlighted that florivory altered specific chemical families rather than the global metabolic fingerprint. These responsive features were predominantly associated with the shikimate and phenylpropanoid pathways, whose associated metabolic intensities mainly increased under aphid and beetle florivory. For instance, florivory significantly affected phenolic acid diversity indices, indicating that florivore attack alters both compositional and functional dimensions of phenolic diversity. Phenolics are widespread plant defense metabolites that deter herbivores and impair digestion, with insect attack inducing changes in their profiles (War et al. 2012; Mostafa et al. 2022). In *Brassica nigra*, florivory by *Pieris brassicae* altered floral phenolic profiles, demonstrating that such chemical defenses also operate in reproductive tissues (Lucas-Barbosa et al. 2016). In line with the framework proposed by Philbin et al. (2022), who showed that herbivores interact with both metabolic composition and structural complexity as distinct dimensions of plant chemical diversity in *Piper* shrub species, our results suggest that florivory may influence phenolic diversity at different levels. In addition to phenolic compounds, a few features assigned to fatty acids also responded to the florivory treatment in *T. vulgare*. Lipid-derived compounds are also involved in herbivore-induced signaling processes. In tomato leaves, simulated wounding caused rapid increases in linolenic acid, which were temporally associated with subsequent jasmonic acid accumulation, supporting the role of lipid release as an early event in defense signaling (Conconi et al. 1996). The fatty acid responses observed in our study may therefore reflect activation of comparable lipid-mediated signaling pathways during floral responses to florivory. Beyond their role in jasmonate biosynthesis, polyunsaturated fatty acids can also be oxidized via the lipoxygenase (LOX) pathway to form oxylipins, including green leaf volatiles that contribute to wound and herbivore signaling (Paul et al. 2022). It is important to consider that the experiment was conducted under field conditions and prior damage by other florivores, particularly by *Olibrus* spp., cannot be excluded, while no aphids were observed on flower stems prior to the treatment.

### Florivore type shapes the direction and magnitude of floral chemical responses

Our results highlight that the floral chemical response was tailored to the insect species attacking *T. vulgare*. (*E*)-β-caryophyllene is commonly reported as an induced volatile following herbivory (Moraes et al. 1998; Abel et al. 2009). In contrast to leaf emission studies, we observed decreased contents of this sesquiterpene in flower heads after beetle florivory, while aphids had no effect. Notably, (*E*)-β-caryophyllene has been reported to act as an antifeedant against the facultative florivore *Metrioptera bicolor* (Junker et al. 2010). Bustos-Segura and Foley (2018) demonstrated that herbivore-induced emissions largely mirrored constitutive terpene pools, suggesting that emissions predominantly originate from stored pools rather than de novo biosynthesis. Measurements of both stored and emitted terpenes would be needed to test how both respond to florivory in *T. vulgare*.

Whereas stored terpene responses suggested only minor florivore-specific effects in our experiment, untargeted metabolomic analyses revealed clearer differentiation between aphid- and beetle-induced metabolic changes. Beetle florivory affected more metabolic features and was mainly associated with increases, whereas aphids predominantly induced decreases. These contrasting patterns may reflect the differences in feeding mode. Phloem-feeding aphids extract nutrients with minimal tissue damage, thereby limiting or even suppressing the activation of damage-dependent defense pathways (Walling 2008). For example, glucosinolate-based defenses require tissue disruption for enzyme–substrate mixing, which phloem feeders can prevent, even reducing glucosinolate levels as shown for *Myzus persicae* infesting *A. thaliana* (Kim and Jander 2007). In contrast, chewing herbivores such as *Psylliodes chrysocephala* induced systemic increases in glucosinolate concentrations in *Brassica napus* (Bartlet et al. 1999). Similar patterns have been reported in other systems, where aphids elicited comparatively weak effects, while chewing herbivores were the primary drivers of floral responses (Chrétien et al. 2018). Likewise, in *Plantago lanceolata*, aphid feeding resulted in only minor metabolic changes, with several features being downregulated in systemic leaves, while chewing herbivores induced upregulation of multiple features in locally damaged leaves (Sutter and Müller 2011). Furthermore, our results showed limited overlap of eatures being significantly induced by aphid and beetle treatments, indicating that different florivores trigger largely distinct metabolic responses, related to the different damage pattern and different signaling pathways that they trigger (Koornneef and Pieterse 2008; Pieterse et al. 2012). Together, these findings support the view that the feeding pattern and mode are key determinants of the magnitude, direction and overall pattern of induced metabolic responses in both vegetative and reproductive plant parts (Sutter and Müller 2011; Pareja et al. 2012; Lucas-Barbosa et al. 2016).

### Chemotype identity modulates floral responses to florivory

Variation in floral metabolic responses was somewhat chemotype-specific. For instance, within the terpenes, caryophyllene oxide decreased after aphid treatment only in the B-Thu chemotype, whereas spathulenol and trans-pinocarveol responded after beetle treatment exclusively in the Keto and AB-Thu chemotypes, respectively. Beyond individual terpenes, the chemotype × treatment interaction in untargeted metabolomic analyses also affected chemodiversity indices linked to fatty acyl glycosides, indicating that chemotype-dependent induction patterns extend across multiple metabolite classes. Moreover, at the level of individual metabolites other than terpenes, significant responses were largely confined to single chemotypes, with minimal overlap among them. These findings support our hypothesis that florivore-induced metabolic responses depend on plant chemotype and are consistent with previous observations in roots and shoots of *T. vulgare* (Newrzella et al. 2026). Comparable patterns have also been described in other species. In wild cotton (*Gossypium hirsutum*), leaf emissions of aldoximes and nitriles, as well as stored concentrations of (*E*)-β-ocimene, varied between chemotypes after caterpillar feeding (Mamin et al. 2025). Similarly, terpene emissions following artificial damage differed among chemotypes of *Melaleuca alternifolia* (Bustos-Segura and Foley 2018). While these studies focused primarily on vegetative tissues, our results demonstrate that chemotype diversity also shapes florivore-induced metabolic responses in flower heads. In conclusion, our study provides novel insights into stem-specific floral metabolic responses that are shaped by both florivore type and plant chemotype. While the overall metabolome remained largely stable, potentially maintaining pollinator attractiveness, distinct changes occurred in specific chemical classes, particularly the shikimate and phenylpropanoid pathways. Aphid infestation was mainly associated with decreases in metabolite levels, whereas beetles mostly induced increases. Future research should further explore how diverse florivores and plant chemotype jointly shape the floral metabolome to better understand the ecological drivers of flower chemistry.

## ACKNOWLEDGEMENTS

We thank Lukas Brokate from Bielefeld University for practical help. We acknowledge support for the publication costs by the Open Access Publication Fund of Bielefeld University.

## AUTHOR CONTRIBUTIONS

SG, CM and TD designed the study. SG performed the experiment and analyzed the data with support from TD. SG generated the figures and wrote the first draft of the manuscript and all the authors contributed to the final version.

## SUPPORTING INFORMATION

Chemical data, metadata and code are available on https://git.nfdi4plants.org/for-3000/Data_Plant_P5_Gaar_2026/-/tree/6001175dac83ab4ac9be2d627a2a4fb87585d536/. Data will be made publicly available upon acceptance of this manuscript.

