## Supplemental figures for "Impacts of different types of florivores on flower metabolomes in the field"

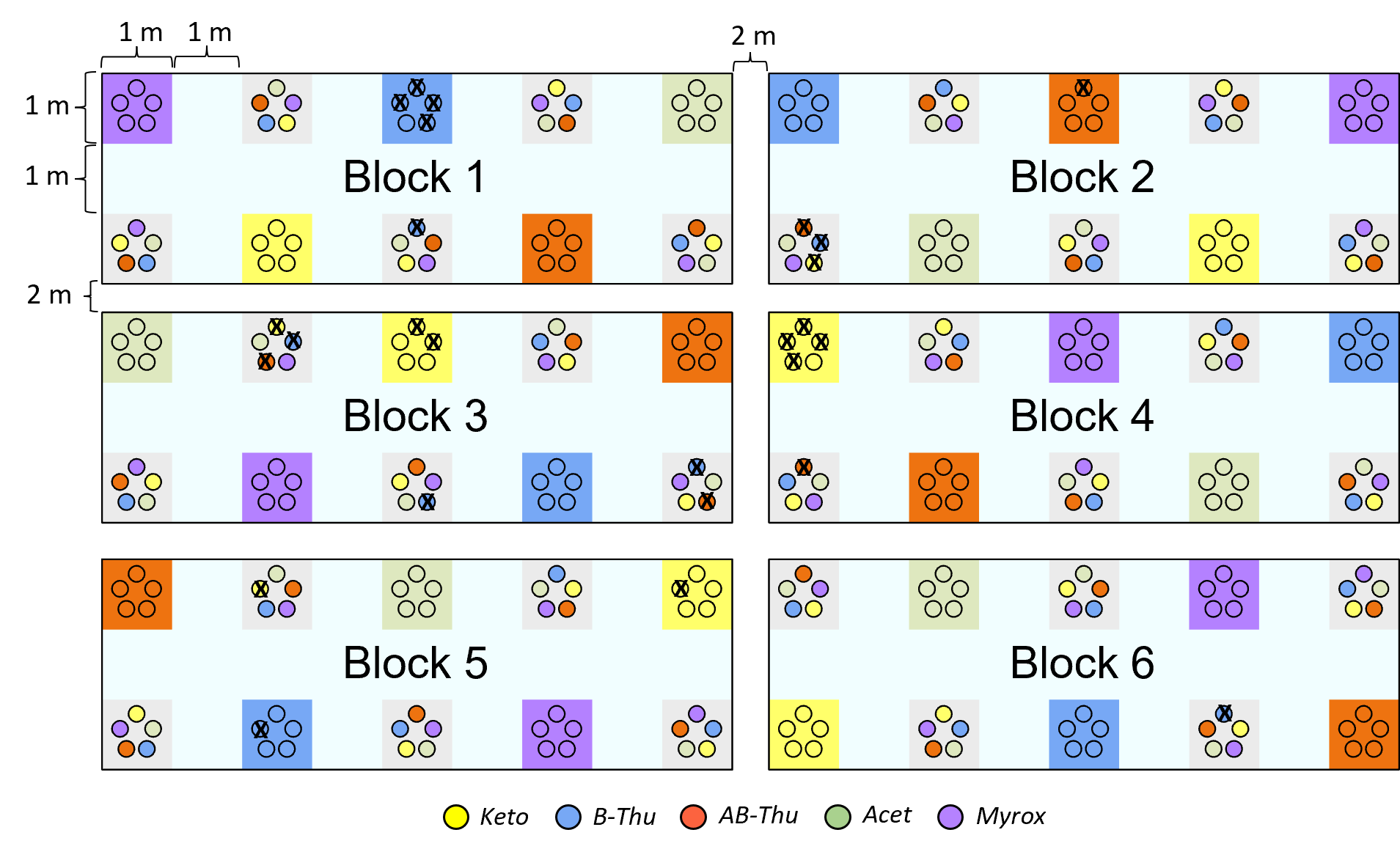

**Fig. S1. Experimental common garden design** (adapted from Ziaja & Müller 2023). *Tanacetum vulgare* plants of five distinct chemotypes were planted in either homogeneous plots (five plants of the same chemotype) or heterogeneous plots (five plants of different chemotypes). Chemotypes are indicated by color: yellow = artemisia ketone (Keto); blue = β-thujone (B-Thu); orange = α-/β-thujone (AB-Thu); green = artemisyl acetate/artemisia ketone/artemisia alcohol (Aacet); and purple = (Z)-myroxide/santolina triene/artemisyl acetate (Myrox). From each plant individual, two clones were represented in the whole field, one in a homogenous plot and one in a heterogenous plot within the same block. Due to waterlogging during the preceding winter, only 30 plants were used in the present experiment in summer 2024 (Keto: *n* = 10; B-Thu: *n* = 11; AB-Thu: *n* = 9). These plants are indicated by an “x”.

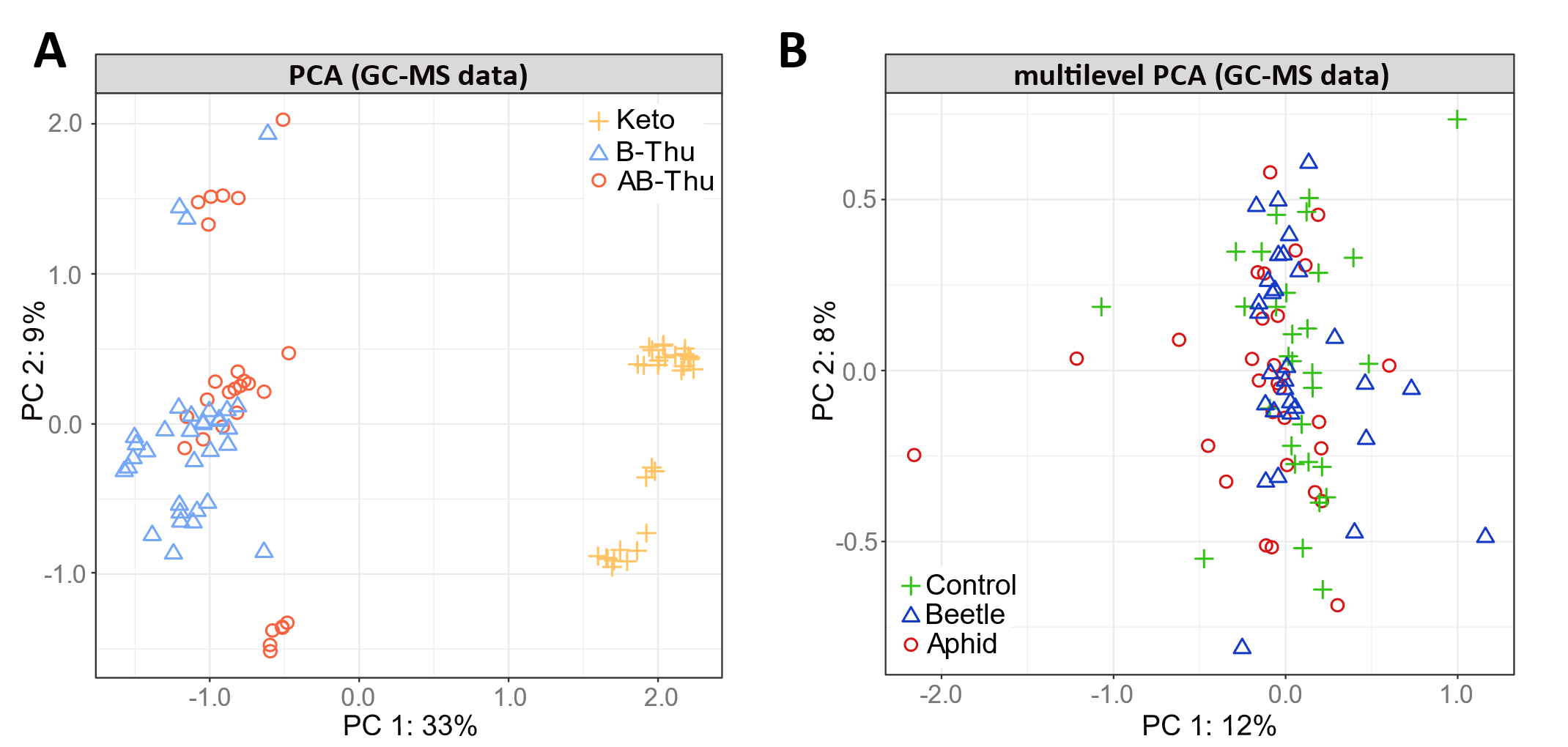

**Fig. S2.** **PCA of stored terpene profiles of *Tanacetum vulgare* flower heads in relation to treatment and chemotype**. **A.** Multilevel PCA score plot of stored terpenes illustrating treatment-related variation in a paired data set. Analysis was performed on normalized GC-MS data (102 terpenes). The unsupervised PCA does not reveal a clear separation between control, beetle-, and aphid-infested samples. Symbols and colors indicate the treatments: green crosses represent uninfested flower head control samples, red circles represent samples infested with aphids (*Macrosiphoniella tanacetaria*) and blue triangles represent samples infested with beetles (*Olibrus* spp.). **B.** PCA score plot of terpene profiles showing separation of chemotypes. Symbols and colors indicate different chemotypes: yellow crosses, artemisia ketone (Keto); orange circles, α-/β-thujone (AB-Thu); light blue triangles, β-thujone (B-Thu).

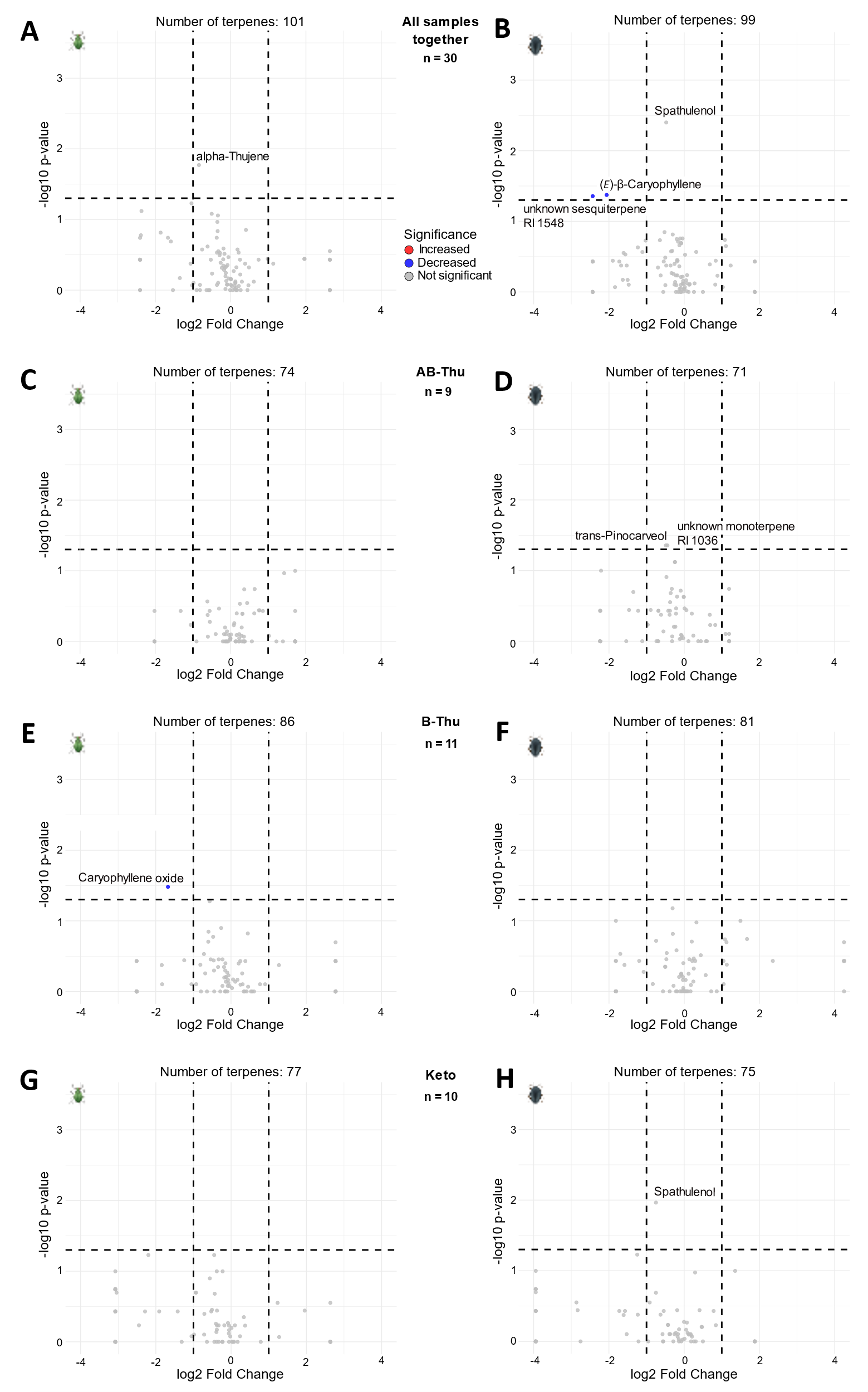

**Fig. S3.** **Species- and chemotype-specific responses in *Tanacetum vulgare f*lower heads of terpenes to aphid and beetle infestation**. Volcano plots showing terpenes significantly modulated by insect infestation, displayed as circles outside the cutoff lines (Wilcoxon test, *P* < 0.05; fold change < 0.5 for decreases and > 2 for increases) based on absolute GC-MS data. Blue circles indicate terpenes with reduced concentrations in infested flower heads compared to controls. For terpenes detected exclusively in one treatment group, fold changes were set to the maximum observed value for decreases or increases within the respective comparison. Numbers shown at the top center indicate the total number of terpenes detected per treatment comparison. Sample sizes (*n*) are given as numbers of paired plants (insect-treated *versus* control). Names of significantly affected terpenes are shown next to the corresponding circles. **A.** All aphid *versus* control samples. **B.** All beetle *versus* control samples. **C.** Aphid *versus* control, α-/β-thujone chemotype (AB-Thu). **D.** Beetle *versus* control, AB-Thu chemotype **E.** Aphid *versus* control, β-thujone chemotype (B-Thu). **F.** Beetle *versus* control, B-Thu chemotype. **G.** Aphid *versus* control, artemisia ketone chemotype (Keto). **H.** Beetle *versus* control, Keto chemotype.

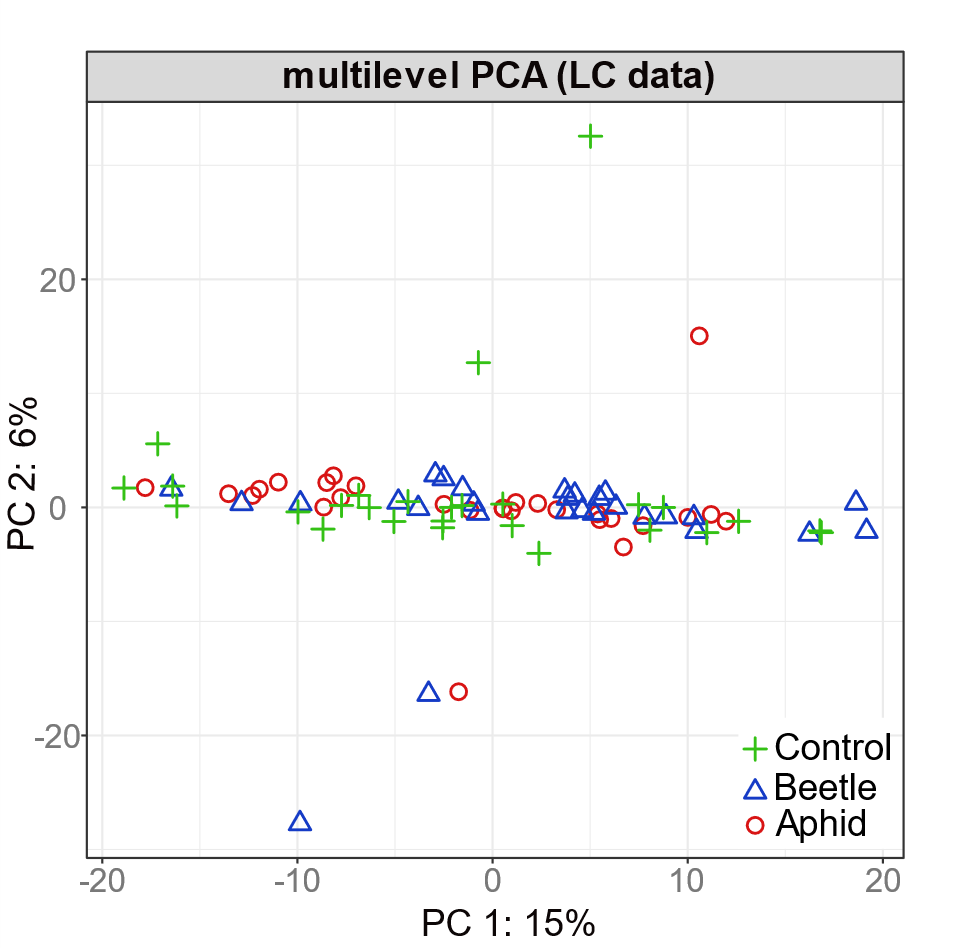

**Fig. S4. PCA of metabolic fingerprints of *Tanacetum vulgare* flower heads in relation to insect treatment**. Multilevel PCA score plot of LC-MS metabolic fingerprints illustrating treatment-related variation in a paired data set. Analysis was performed on normalized data comprising 3,731 features. The unsupervised PCA does not reveal a clear separation between control, beetle-, and aphid-infested samples. Symbols and colors indicate the treatments: green crosses represent uninfested flower head control samples, red circles represent samples infested with aphids (*Macrosiphoniella tanacetaria*) and blue triangles represent samples infested with beetles (*Olibrus* spp.).

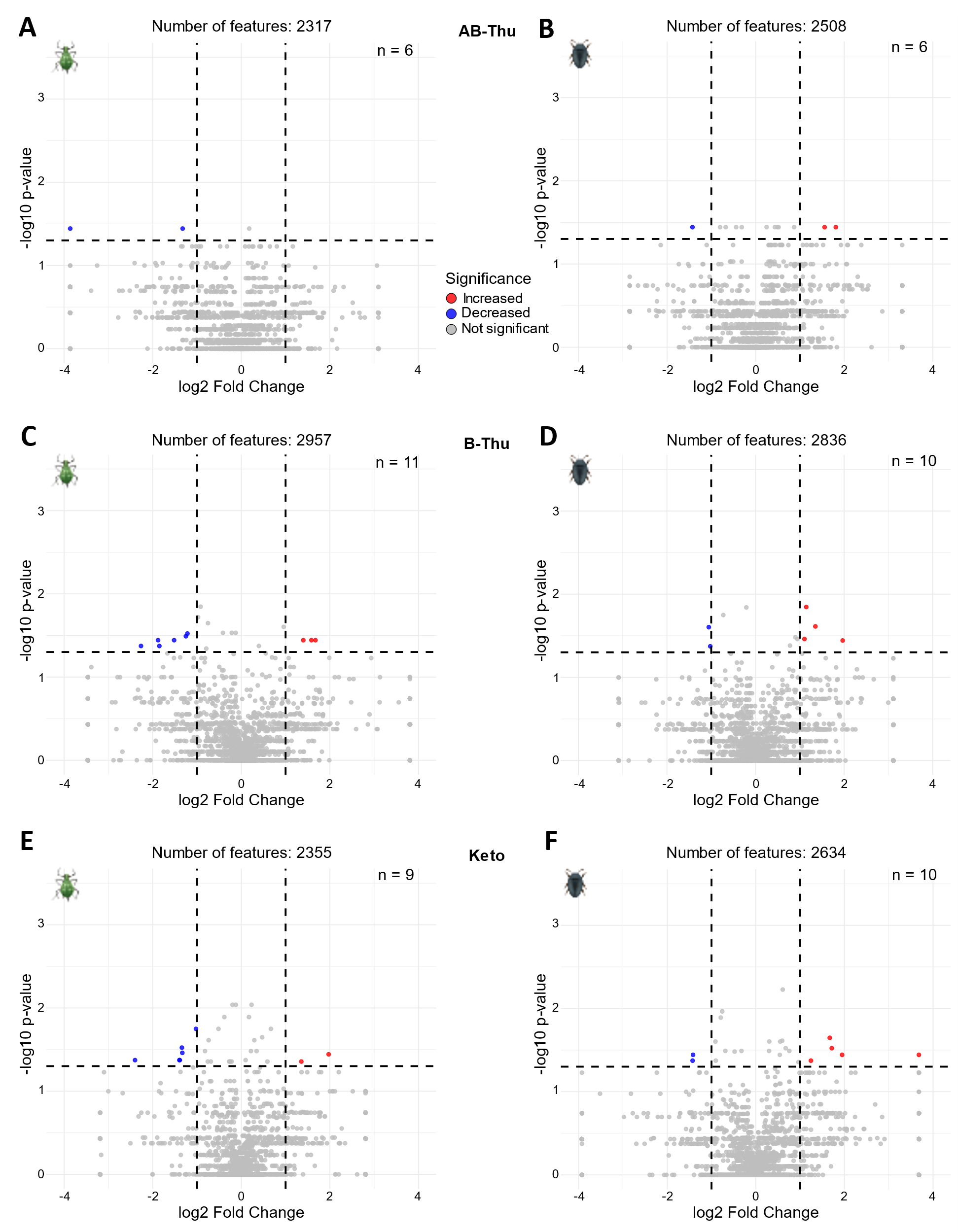

**Fig. S5. Species- and chemotype-specific metabolic responses of *Tanacetum vulgare* flower heads to aphid and beetle infestation**. Volcano plots showing metabolites significantly modulated by insect infestation, displayed as circles outside the cutoff lines (Wilcoxon test, *P* < 0.05; fold change < 0.5 for decreases and > 2 for increases) based on absolute LC-MS data. Blue circles indicate metabolites with reduced concentrations in infested flower heads compared to controls, while red circles indicate increased concentrations. For metabolites detected exclusively in one treatment group, fold changes were set to the maximum observed value for decreases or increases within the respective comparison. Numbers shown at the top center indicate the total number of terpenes detected per treatment comparison. Sample sizes (*n*) are given as numbers of paired plants (insect-treated *versus* control). **A.** Aphid *versus* control, α-/β-thujone chemotype (AB-Thu). **B.** Beetle *versus* control, AB-Thu chemotype. **C.** Aphid *versus* control, β-thujone chemotype (B-Thu). **D.** Beetle *versus* control, B-Thu chemotype **E.** Aphid *versus* control, artemisia ketone chemotype (Keto). **F.** Beetle *versus* control, Keto chemotype.
